# Multimodal memory components and their long-term dynamics identified in cortical layers II/III but not layer Vb

**DOI:** 10.1101/665539

**Authors:** Dong Li, Guangyu Wang, Hong Xie, Yi Hu, Ji-Song Guan, Claus C. Hilgetag

## Abstract

Activity patterns of cerebral cortical regions represent the present environment in which animals receive multi-modal inputs. They are also shaped by the history of previous activity that reflects learned information on past multimodal exposures. We studied the long-term dynamics of cortical activity patterns during the formation of multimodal memories by analysing *in vivo* high-resolution 2-photon mouse brain imaging of Immediate Early Gene expression, resolved by cortical layers. Strikingly, in layers II/III, the patterns showed similar dynamics across functional distinct cortical areas and the consistency of dynamic patterns lasts for one to several days. In contrast, in layer Vb, the activity dynamics varied across functional distinct areas, and the present activities are sensitive to the previous activities at different time depending on the cortical locations, indicating that the information stored in the cortex at different time points is distributed across different cortical areas. These results suggest different roles of layer II/III and layer Vb neurons in the long-term multimodal perception of the environment.

## Introduction

The brain can represent, integrate and remember information from more than one sensory modality [1; 2; 3; 4; 5]. This cross-modal integration is structured such that items can be represented both as a whole as well as a set of cross-model details. In a complex environment, the learning of these integrated representations is a difficult task requiring repeated exposures to the multi-sensory stimuli. Moreover, the learning mechanisms need to address plasticity-stability trade-offs, by forming relevant new cross-modal associations while ignoring and forgetting irrelevant associations and preserving prior memories. As a result, the formation of cross-modal memories becomes a long-term dynamic process. Understanding the long-term dynamics of cortical memory representation in multimodal environment is not only a worthwhile topic by itself in brain research, but also significant for inspiring the enhancement of cross-model learning abilities of artificial brains [6].

The cerebral cortex of the mammalian brain, which is parcellated into a multitude of structurally and functionally specific, layered areas, is believed to be involved in higher-order brain functions, including multisensory perception [1]. Substantial evidence suggests that the cerebral cortex has both area-specific and layer-specific functions in the processes of learning and memory. For example, Phoka et al [7] found increased neural activity and concomitant ensemble firing patterns in mouse somatosensory cortex, specifically layers IV and Vb, sustained for more than twenty minutes after multi-whisker, spatiotemporally rich stimulation of the vibrissae. Kitamura et al [8] pointed out that contextual fear memory can be quickly produced at the onset of learning in prefrontal cortex. Xie et al [9] discovered memory trace neurons in layer II/III of various areas of mouse cortex. Wang et al [10] demonstrated that the cross-modal integration of visual and somatosensory inputs evoked specific neural responses in particular cortical areas, such as the primary visual cortex and retrosplenial cortex. Sellers et al [11; 12]demonstrated that anesthetics could selectively alter spontaneous activity as a function of cortical layer and suppress both multimodal interactions in primary visual cortex and sensory responses in prefrontal cortex. Despite these extensive observations, however, it remains unclear whether and how the long-term dynamics of cortical memory representations are cortical area- and layer-specific.

In this study, we investigate the long-term dynamics of cortical area and layer-distributed cellular activity patterns during the formation of cross-model memories by analysing *in vivo* high-resolution 2-photon imaging data from BAC-EGR-1-EGFP mouse brains in multimodal environments. On each day, animals were put into one type of environment, receiving multimodal inputs. Several cortical locations from various regions of each subject were monitored, and within each location the neural activity patterns were represented by the firing rates of 6,000 to 15,000 neurons, across multiple cortical layers. During memory formation, the activity patterns of a particular day could be related to those on previous days, which was analysed using a prediction algorithm by a gradient boosting decision tree implemented in the LightGBM Python-package [13]. We show that the long-term memory-related cortical dynamics are significantly layer specific. In layers II/III, the dynamics are similar across different types of cortical areas and different hemispheres, and the neural activities have an unspecific memory effect, that is they are always more sensitive to the recent history of one to several days than those of longer time ago, even if the more recent memories belong to different environments from the present one. In layer Vb, the dynamics vary among cortical locations, as the information stored in this laminar compartment at different previous time points is distributed across different cortical areas. Those results, therefore, suggest different roles of layer II/III and layer Vb neurons in the multimodal perception of the environment.

## Methods

### Animal experiments

We analysed data from four mice. The used mouse strain was BAC-EGR-1-EGFP (Tg(Egr1-EGFP)GO90Gsat/Mmucd from the Gensat project, distributed by Jackson Laboratories. Animal care was in accordance with the institutional guidelines of Tsinghua University, and the whole experimental protocol was also approved by Tsinghua University. Imaging and data acquisition procedures were previously described by Xie et al [9]. Specifically, mice were 3-5 months old, and received cranial window implantation; recording began one month later. To implant the cranial window, the animal was immobilized in custom-built stage-mounted ear bars and a nosepiece, similar to a stereotaxic apparatus. A 1.5 cm incision was made between the ears, and the scalp was reflected to expose the skull. One circular craniotomy (6-7 mm diameter) was made using a high-speed drill and a dissecting microscope for gross visualization. A glass-made coverslip was attached to the skull. For surgeries and observations, mice were anesthetized with 1.5% isoflurane. EGFP fluorescent intensity (FI) was imaged with an Olympus Fluoview 1200MPE with pre-chirp optics and a fast AOM mounted on an Olympus BX61WI upright microscope, coupled with a 2 mm working distance, 25x water immersion lens (numerical aperture 1.05).

We employed several types of environments for the animals. In principle, they were all multimodal environments, but of different complexity in terms of the sensory modalities. Home Cage was considered the default, where, although the animals could see and touch the cage, as well as smell their own smells, they adapted to this environment and were closely familiar with the sensory inputs. Therefore, the visual, somatosensory and olfactory inputs in the Home Cage environment were all considered as weak, and this multimodal environment was considered as the simplest one compared to all others. More complexity was created by introducing stronger and specific inputs of certain modalities. To this end, we used another three boxes, labelled as contexts A, B, and C, which comprised different shapes, colors, materials of the floors and combinations of different smells, so that animals received strong and specific visual, somatosensory, and olfactory inputs. In addition, in box C, we also employed strong light and sound stimuli. When an animal was put into one of the boxes, it could experience three types of situations. Training A, B, or C meant the animal received foot shocks that are strong enough to lead to freezing behaviors, as part of conditioning for learning. At the same time, the foot shock could also be considered as a very strong and special somatosensory (nociceptive) input by itself. When the animal did not receive the foot shock, we labeled the boxes as Context A, B, or C if before training or as Retrieval A, B, or C after training, respectively. Training C had the largest complexity in terms of sensory modalities compared to the others, and interestingly, in the pre- and post-training phases, the animals displayed different behaviors, that is the freezing in Retrieval A, B, or C but not in Context A, B, or C [9], but we assumed the provided sensory information was identical between the Context and Retrieval environments. Several other environments were also employed, which were more complex than the Home Cage, but simpler than those mentioned before. Enriched Environments and Tunnel were two boxes where animals could receive strong visual and somatosensory inputs. Another two simple environments where employed where the animals only received visual inputs of vertical or horizontal stripes.

Illustrations of the different environment are given in Fig. 1A and the sensory modalities of all environments are summarized in Table 1. The environments that the four mice experienced respectively are summarized in Table 2.

**Figure 1.**
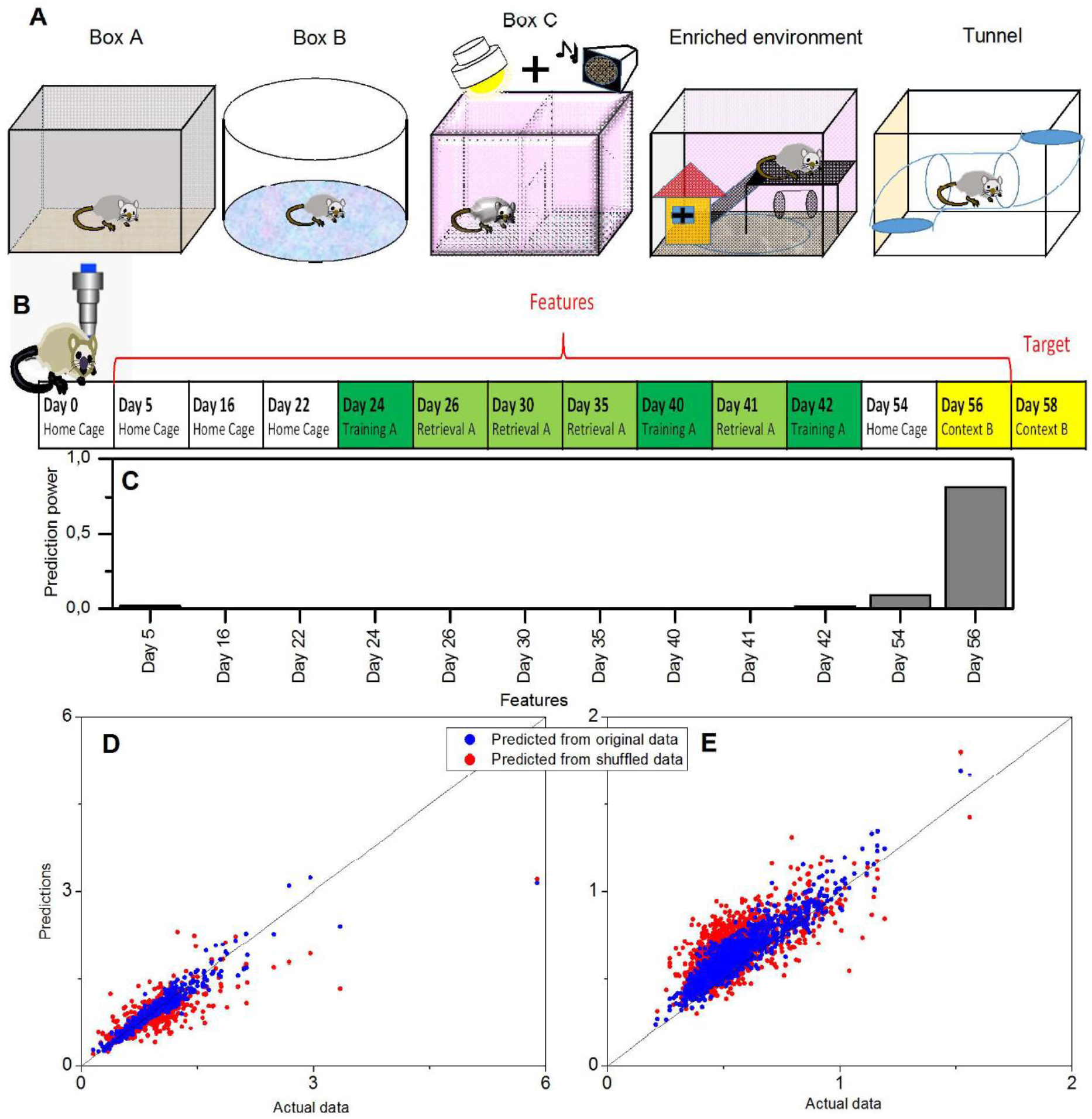
Example of a prediction of memory trace activities. (A) Illustration of different environments employed in this paper. (B) The model was trained on layers II/III in the cortical location A1 (VISp, left) of animal Ma. Neural activities on Day 58 (context B) were used as the target, and the data on twelve previous days were used as the features. (C) Prediction powers of the twelve days in features. (D) Prediction performance of the model on other layers II/III neurons within the same location, and (E) prediction performance of the same model on all layer Vb neurons within the same location, where blue dots indicated the prediction from the original data [R^2^=0.82 in panel (D) and R^2^=0.57 in panel (E)] and red dots from the shuffled data [R^2^=0.61 in panel (D) and R^2^=-0.13 in panel (E)].

**Table 1.**
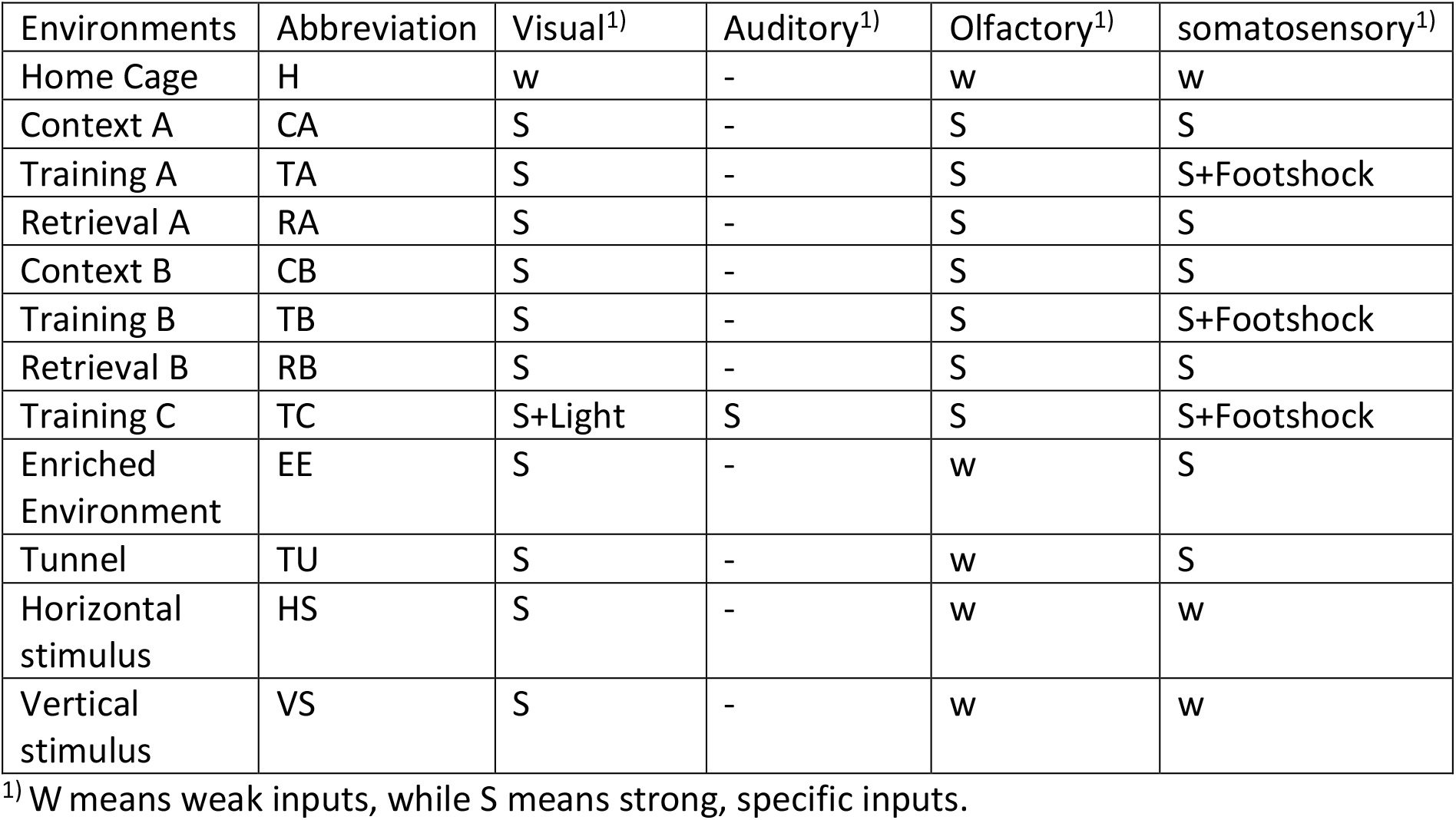
List of the used multimodal environments

**Table 2.** Summery of the subjects

| Mouse | Scanned length (day)<sup>1)</sup> | Scanned times | Environments | Selected Locations | Cortical areas covered<sup>2)</sup> | Minimal number of neurons in layers II/III in each location | Maximal number of neurons in layers II/III in each location | Minimal number of neurons in layer Vb in each location | Maximal number of neurons in layer Vb in each location |
| --- | --- | --- | --- | --- | --- | --- | --- | --- | --- |
| Ma | 131 | 32 | Home cage<br>Training A<br>Retrieval A<br>Context B<br>Training C<br>Enriched Environment<br>Tunnel | 7 | PTLp (L)<br>SSp (L)<br>VISp (L)<br>RSC (R)<br>VISp (R) | 3683 | 6158 | 311 | 1593 |
| Mb | 52 | 22 | Home Cage<br>Training B<br>Retrieval B<br>Context A | 8 | RSC (L)<br>VISam (L)<br>VISp (L)<br>Motor (R)<br>RSC (R)<br>VISam (R) | 2041 | 4863 | 399 | 1104 |
| Mc | 61 | 12 | Home cage<br>Training A<br>Retrieval A<br>Context B | 6 | PTLp (L)<br>RSC (L)<br>VISam (L)<br>PTLp (R)<br>VISam (R) | 842 | 3775 | 116 | 549 |
| Md | 55 | 26 | Home cage<br>Horizontal stimulus<br>Vertical stimulus<br>Enriched Environment | 3 | Motor (R)<br>SSp (R) | 2821 | 5492 | 50 | 1400 |

1) Calculated from the first to the last day of scanning, and signed as Day 0, Day 1, etc.
2) L and R mean the left and right hemispheres, respectively.

### Data selection

For each mouse, typically 10 to 20 cortical locations were monitored, but we only selected those ones that could been scanned at least to a depth of layer Vb for all days of scanning. As a result, we selected 7, 8, 6, and 3 locations for those four mice, respectively, which covered motor, posterior parietal (PTLp), retrosplenial cortex (RSC), primary somatosensory (SSp), anterior medial visual (VISam), and primary visual (VISp) cortical areas on both left and right hemispheres. The neuron positions in the images were automatically detected, as described in detail by Xie et al [9]. If a neuron was missed in the detection for more than three days, the neuron would be excluded from the analysis, and if a neuron was missed in the detection for at most three days, its missed activity values were filled as the median value of all the other neurons on that day. The area types and laminar compartments were manually annotated based on their cytoarchitecture by one expert (G.W.) and approved by all other experimental experts among the authors (H.X., Y.H., and J.-S. G.). In this study we focused our analysis on the activities in layers II/III and layer Vb. A summary of the data available for the analysed four animals is given in Table 2.

### LightGBM prediction approach

We analysed the long-term dynamics of cortical memory representations, as a regression problem, by predicting the activity pattern on a certain day based on the past history of activity patterns. Practically, we used the gradient boosting decision tree implemented in LightGBM [13] Python-package

For each prediction, we needed to select training, validation and test data. Once the activities on a certain day were selected as the target, their values in the training and validation data sets were used as the labels. The values in the test data were not used in the prediction process, but only as ground truth to evaluate the prediction performance. Features were the activities on the previous days. The parameters used in the LightGBM prediction are listed in Table 3.

**Table 3.** Parameters for LightGBM prediction.

| Num_leaves | objective | min_data_in_leaf | Learning_rate | feature_fraction | bagging_fraction | bagging_freq | metric | num_threads |
| --- | --- | --- | --- | --- | --- | --- | --- | --- |
| 10 | regression | 1 | 0.05 | 0.93 | 0.93 | 1 | l2 | 4 |

Since we used ‘12’ for the parameter “metric” in the evaluation process (which means that the mean square error was the target to be optimised in the process of the regression), we calculated the mean square error δ between the prediction results and the ground truth as accuracy estimate. To generate controls, we shuffled the data on the feature days for each neuron.

### Cross-location prediction

For each animal, we selected one specific laminar compartment ∧ (∧ was either in layers II/III or layer Vb). One model was trained by using the training and validation data from one cortical location i_∧_, and predictions were subsequently performed by using the test data from a different location j_∧_ in the same laminar compartment. In this part, the target was always selected as the data on the last day when the mouse’s brain was scanned, and the features were the data on all the previous days that were available, excluding Day 0, in total from 10 to 30 days (see Table 4). To make all pairs of predictions comparable, in this part, for each mouse we needed to select the data sizes of training, validation and test data, respectively, to be always identical for every model. To this end, for each mouse, we first found the minimal number of neurons of every location in layers II/III or layer Vb in the data set, which turned out to be 311, 399, 116, and 50 neurons, respectively (Table 2). This number was the size of test data for each mouse, and the size of training and validation data were 90% and 10% of these numbers, respectively, as seen in Table 4. With those fixed numbers, the data sampling was random, and the validation and the training data sets never had overlaps.

**Table 4.** Summary of cross-location prediction

| Mouse | Number of days in features | Data points in training / validation / test set | Total pairs of comparison | Number of worse performance (significant ones)<sup>1)</sup> | Number of better performance (significant ones)<sup>1)</sup> | Worst difference in performance<sup>1)</sup> | Best difference in performance<sup>1)</sup> |
| --- | --- | --- | --- | --- | --- | --- | --- |
| Ma | 30 | 297/33/311 | 42 | 38 (24) | 4 (2) | -0.498 | 0.124 |
| Mb | 20 | 359/39/399 | 56 | 38 (25) | 18 (5) | -0.351 | 0.234 |
| Mc | 10 | 104/11/116 | 30 | 21 (7) | 9 (3) | -0.192 | 0.126 |
| Md | 24 | 45/5/50 | 6 | 4 (4) | 2 (0) | -0.434 | 0.023 |
| sum | - | - | 134 | 101 (60) | 33 (10) | - | - |

1) Compared the performance in layer Vb to layers II/III

To evaluate the prediction results, we not only calculated the square error δ(i_∧_, j_∧_), but also shuffled the data on the feature days in the test data sets 20 times for the comparison of each pair of training-test locations, and predicted the target each time, so as to obtain another 20 predicted results. The control square error δ_s_(i_∧_, j_∧_) was calculated by using the average of the 20 predicted results from the shuffled data. The relative error was then calculated as δ_r_(i_∧_, j_∧_)=δ(i_∧_, j_∧_)/δ_s_(i_∧_, j_∧_). We defined the prediction quality measurement k as k(i_∧_, j_∧_)=exp(-δ_r_(i_∧_, j_∧_)), and the matrix M_k∧_, whose off-diagonal entry at the i^th^ row and j^th^ columns was k(i_∧_, j_∧_) and diagonal elements were all empty. M_k∧_ was, therefore, able to reflect how the memory-dependent dynamics of the neural populations from the testing location were similar to the training location.

We repeated these predictions and evaluations 10 times, so as to obtain 10 M_k∧_. The differences in prediction performances for layers Vb and II/III could be demonstrated in two ways. In the first instance, we averaged all the 10 M_k∧_ for each layer compartment, to obtain 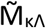, and 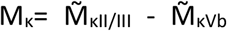, and finally used the matrix M_s_=(M_k_+ M_k_^T^)/2 to demonstrate the difference. If one entry was 0, it means the predictions for layer Vb and layer II/III had the same performance in the corresponding pair of locations, and the values larger (or smaller) than 0 mean prediction in layers II/III (or layers Vb) had better performance. In the second instance, we directly compared the difference of the 10 values between k(i_Vb_, j_Vb_) and k(i_II/III_, j_II/III_), to search for the significant difference (p<0.01, t-test with Bonferroni correction).

### Intra-location prediction

Intra-location prediction was basically performed in the same way as cross-location prediction. The only difference was that, since the test data set came from the same population as the training and validation data sets, it was necessary to make sure that those data did not have overlaps. To this end, we equally divided the data in each location and laminar compartment into 10 groups. For each prediction, we sampled one group of neurons, and randomly sampled 50 neurons from this group as the validation data, sampled another two groups of neurons, and randomly sampled 100 neurons from these two groups as the test data, and randomly sampled 350 neurons from the left groups as the training data. Locations whose layers II/III or layer Vb did not contain at least 500 neurons would be excluded from this part of analysis.

### Prediction power

Once a model Mod^(r)^ was trained, and we signed the set of feature days as S^(r)^, LightGBM could return the total gains of splits for each feature G_D_^(r)^, where D indicates the feature day used in this model. We therefore directly used the gain normalized by their summation, i.e. 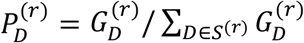 to indicate the prediction power of the feature day D in model Mod^(r)^. Because the prediction power was the property within a model itself, insensitive to its performance with test data, and the measurement was a value normalized within the model, for each model we used a big part (90%) of the neurons within the population (80% as training data and 10% as validation data).

### Repeat environment prediction

In this part, we used four features to predict the target activities. Day 0 was always excluded from the analysis, and the data of the next three scanning time points (labelled as Day S1, Day S_2_ and Day S_3_) were always included in the features, in order to generate controls to evaluate the prediction performance. However, in order to eliminate the predictive effects from those three days that could be different among the situations which we were going to compare, we shuffled the neurons on each of those three days. For each animal, from Day S_4_ on, we were looking for the next scanning day on which the mouse was put into a repeated environment for the first time, and included this pair of repeated environments into the analysis, except for that between those days, the mouse used to be put into the same box, even though in a different environment. For instance, in the sequence consisting of Retrieval A (Day S_n_), Home Cage (Day S_n+1_), and Retrieval A (Day S_n+2_), the pair of Day S_n_ and Day S_n+2_, which has the environment-repeat interval *Inv*=S_n+2_-S_n_, would be included in the analysis, but in the sequence consist of Retrieval A (Day S_n_), Training A (Day S_n+1_), and Retrieval A (Day S_n+2_), the pair of Day S_n_ and Day S_n+2_ would be excluded. For each selected pair, we used the data on the earlier day together with the aforementioned shuffled data on Day S_1_, Day S_2_ and Day S_3_ to predict the activities on the later day, which resulted in a mean square error δ(i_∧_) in location i_∧_ (in layer compartment ∧), and we shuffled the days in the test data, resulting in δ_s_(i_∧_), so eventually we obtained δ_r_(i_∧_)=δ(i_∧_,)/δ_s_(i_∧_), which measured the performance of this prediction, where smaller δ_r_(i_∧_) indicates better prediction. We repeated the prediction 100 times within each location i_∧_, and obtained the averaged value <δ_r_(i_∧_)>, where <·> stands for the average over trials. Within each location i_∧_, we still randomly divided the neurons into ten groups, and for each prediction, we randomly selected four groups (40% of the data) as the training data, one group (10% of the data) as the validation data, and left the other five groups (50% of the data) as the test data.

We calculated the average of <δ_r_(i_∧_)> among all the locations of the mouse, to get the mean value 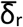 and the standard deviation σ(δ_r_), so that we could analyse their dependence on the environment-repeat interval *Inv*, simply by using lining fitting 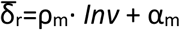 and σ(δ_r_)=ρ_s_· *Inv* + α_s_, respectively. To analyse their dependence on the multimodal environments, we selected the two most often repeated environments for each mouse (eventually 5 to 8 repeating times), and compared 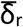 and σ(δ_r_) over all the repeats between those two environments for twice. First, we made the comparison by using the original values, and afterwards in order to eliminate the influence of the different environment-repeat interval *Inv* as much as possible, we made the comparison again by using a kind of modified values, which equalled to the original values minus *Inv* times the fitted slopes, namely 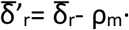 *Inv*, and σ′(δ_r_) = σ(δ_r_) – ρ_s_· *Inv* Respectively.

Subject Mc was excluded from this part of the analysis, because it only experienced very few environment repeats.

## Results

### Predictability

Activities of cortical neurons could indeed be predicted by using gradient boosting decision tree, taking their past activities as features and already knowing some of the activities at the target day as the training labels (example shown in Fig. 1). Although prediction performance varied, the prediction generally worked, even when trying to predict the activities in a different laminar compartment (Fig. 1D) or in a different cortical location (Figs. 2A, 2B, and 2C) from that by which the model was trained.

**Figure 2.**
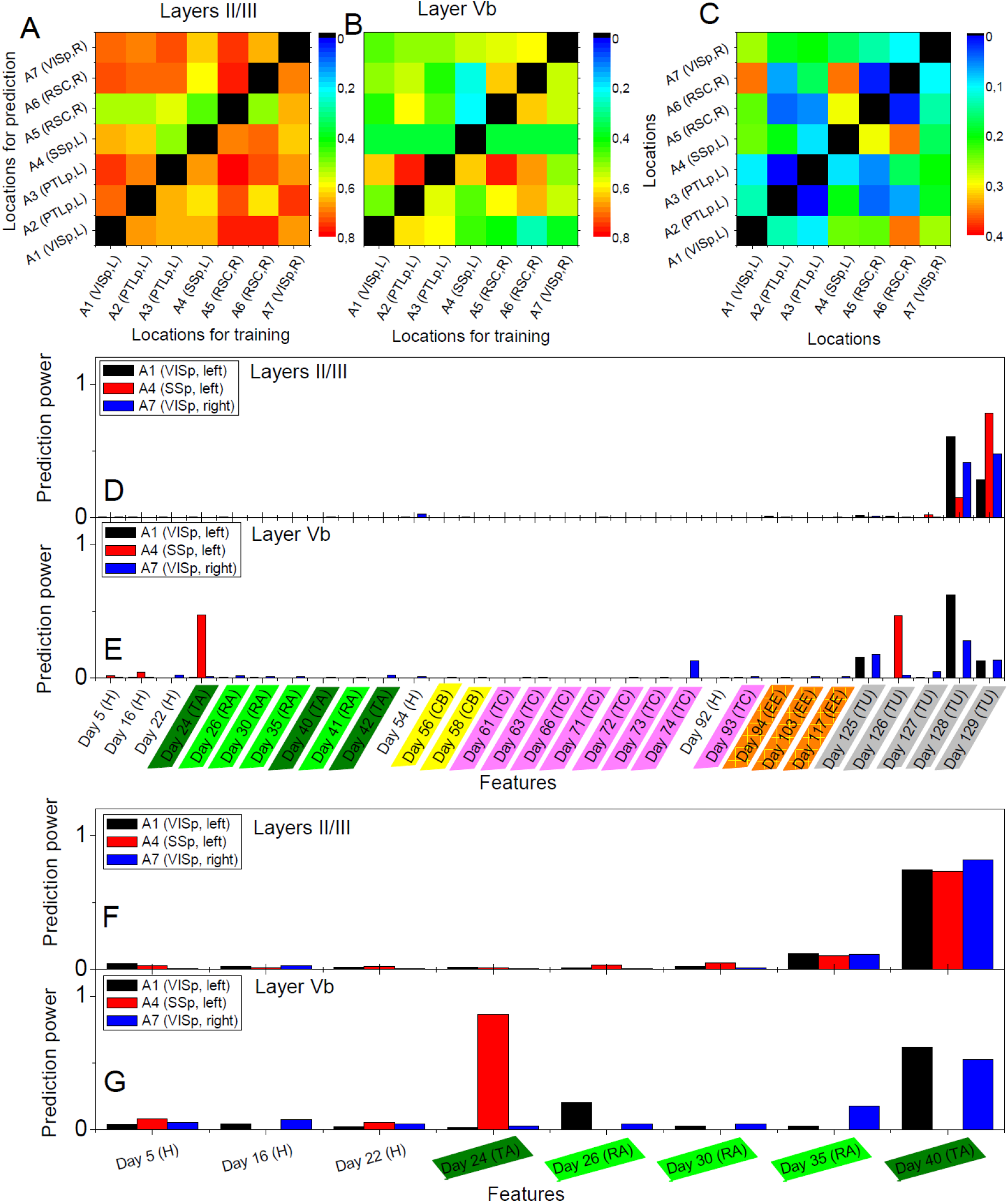
Cross-location prediction performance 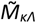 in layers II/III (A) and in layer Vb (B), and their relative difference M_s_ (C), by using the data from animal Ma. Diagonal elements do not have values. Prediction power distributions of layer II/III model (D) and layer Vb model (E) trained in cortical location A1 (left VISp), A4 (left SSp), and A7 (right VISp), when the neural activities on Day 130 (Tunnel) was used as the target and all previous days in the data set as the features. (F) and (G) show the prediction power distributions of layer II/III model and layer Vb model, respectively, when the neural activities on Day 41 (Retrieval A) was used as the target and 8 previous days in the data set as the features. Abbreviations – H: Home cage; TA: Training A; RA: Retrieval A; CB: Context B; TC: Training C; EE: Enriched environment; and TU: Tunnel.

### Cross-location and intra-location predictions

The performance of cross-location prediction was significantly layer-specific. In layers II/III, any model trained from one cortical location could well predict the neural activities in other cortical locations, no matter whether they belonged to the same type of cortical area or the same hemisphere (Fig. 2A). In comparison, cross-location prediction performed much worse in layer Vb (Figs. 2B, 2C and Table 4). Specifically, when we compared the different prediction performances in layer Vb to layers II/III of each pair of training-test locations, for all four animals among all the 134 pairs, we obtained 101 worse performances in layer Vb compared to layers II/III (in terms of the averaged value κ), out of which 60 were significant (p<0.01, t-test with Bonferroni correction within each animal), whereas we had only 33 better performances in layer Vb, out of which only 10 were significant (Table 4).

Intra-location prediction showed the same bias, that is, it performed worse in layer Vb than layers II/III, but the difference was much less significant than cross-location prediction (compared Table 5 to Table 4). Specifically, among all the 13 comparisons, there was only one result showing significant difference.

**Table 5.**
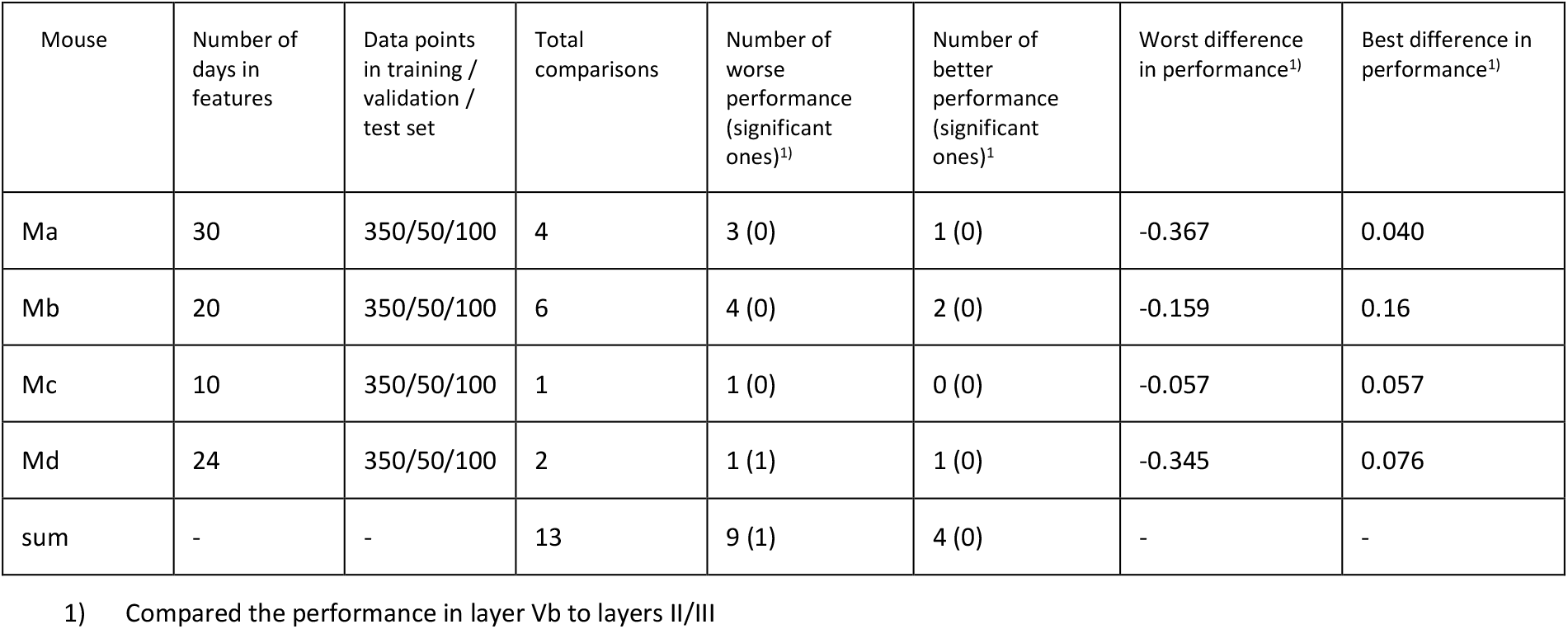
Summary of intra-location prediction

Furthermore, we found that in the cross-location prediction, the large differences in performance tended to appear to the pair of locations with different types of cortical areas (see for example locations A1 and A4 in Fig. 2C, which were in left VISp and left SSp, respectively) or different hemispheres (see for example locations A1 and A7 in Fig. 2C, which were in left VISp and right VISp, respectively).

The analysis of the prediction powers of the days in history helped us obtain deeper insights into the differential performances of layers Vb and II/III. Taking the models trained in A1 (left VISp), A4 (left SSp), and A7 (right VISp) for example, the distributions of the prediction powers for the models in layers II/III were very similar (Fig. 2D). Specifically, most powerful predictors were those on the most recent days (like Day 128 and Day 129 when the targets were on Day 130). In layers Vb, the prediction power had significantly different distributions among the models trained in those three locations (Fig. 2e), where for A1 and A7, three days (Day 125, Day 128 and Day 129) with the same environment as the target day (Tunnel) had high prediction powers and only for A7 another day (Day 74) also had high prediction power, whereas for A4, two days (Day 24, Training A and Day 126, Tunnel) had significantly high prediction powers. Even if we used the data within a short duration in those three locations to train models [for example, Day 41 (Retrieval A) as the target day and all previous days as features], we can still find those different patterns of the prediction power distributions between layers II/III and layer Vb. In layers II/III, the distributions were still very similar (Fig. 2F), but in layer Vb, the distributions were widely different (Fig. 2G).

### Repeat environment prediction

For each mouse, 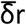 in layers II/III was always more sensitive to environment-repeat interval *Inv* compared to layer Vb, reflected by the bigger slopes ρ_m_, or bigger *R*^2^ values of the lining fitting results, or both (the first column of Fig. 3). σ(δr) did not have very strong interrelation with *Inv*, (the second column of Fig. 3). In the comparisons of 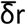 and σ(δr) with the original values between the most often repeated environments, we only found one result which had statistical significance (p<0.5), which was the σ(δr) in layer Vb of mouse Ma between Training C and Tunnel. After modifying the values, the significance did not change too much (p is still smaller than 0.1). Other comparisons that had small p values (<0.1) included 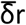 in layers II/III of mouse Mb between Context A and Retrieval B (p>0.1 after the modification), 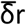 on layers II/III of mouse Md between Enriched Environment and Home Cage (p>0.1 after the modification), σ(δr) on layers II/III of mouse Md between Enriched Environment and Home Cage (p<0.05 after the modification), and σ(δr) on layer Vb of mouse Md between Enriched Environment and Home Cage (p>0.1 after the modification). In addition, σ(δr) in layers II/III of mouse Ma between Training C and Tunnel did have a small p value (p>0.01), but it became smaller than 0.01 after modification.

**Figure 3.**
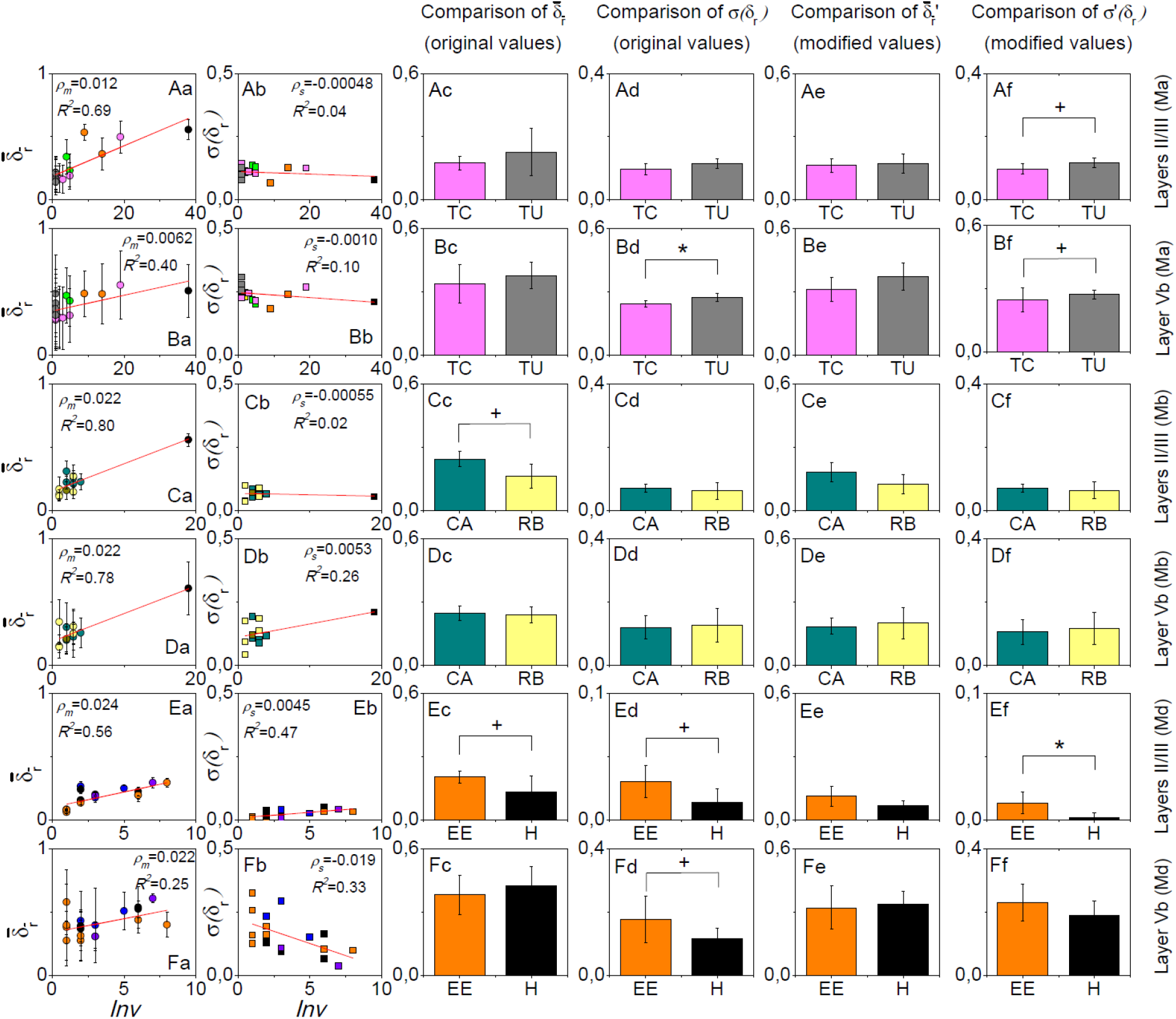
Results of the environment-repeat prediction. Rows from top to bottom are layers II/III of mouse Ma, layer Vb of mouse Ma, layers II/III of mouse Mb, layer Vb of mouse Mb, layers II/III of mouse Md, and layer Vb of mouse Md. The first column is the dependence of 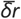 on the environment-repeat interval Inv, where error bars in fact indicate the standard deviation σ(δr), and the red lines are the linear fitting results. The second column shows the dependence of σ(δr) on Inv. The third to the sixth columns show the comparisons of 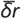 and σ(δr) with the original and modified data, respectively, between the most often repeated environments that each mouse experienced. Colours in the figure are used to discriminate environment types. +: p<0.1 and *: p<0.05.

## Discussion

### Interpretation of the prediction

Although cortical activity patterns in the context of learning and memory appear very complex, they are not purely random. Rather, they are sensitive to outside stimuli as well as their own histories[14]. The prediction approach employed in this work indeed follows such a hypothesis, that cortical neurons can represent long-term memories in multimodal environments, so as to have long-term memory-dependent dynamics. If a model trained within one neural population can also successfully predict the neural activities in another population, it means that within the considered history period, those two populations have similar memory-dependent dynamics. Moreover, the features with high prediction powers indicate the day when the fresh information in the history that is useful for forming the present activity pattern starts to encode in the neural populations. However, the days of the features with very low prediction powers do not necessarily mean that their activities do not correlate with the activities on the target day. Another possibility may be that they do not encode additional useful information for predicting the neural activities on the target day, on top of the days of higher prediction powers.

As a result, we show that within the same cortical location and same laminar compartment, the neurons indeed have similar long-term memory-dependent dynamics. Even across layers, or across areas, the neurons may still have certain similarities in this long-term memory-dependent dynamics, but the similarities vary from case to case.

### Comparison between layers II/III and layer Vb

In this study, we focussed on layers II/III and layer Vb for several reasons. First of all, both layers II/III and deep layers have been shown to play important roles in learning and memory in previous studies [9; 15; 16]. Secondly, the data under study have good quality in multiple locations scanned until layer Vb, and both layers II/III and layer Vb are easy to manually discriminate in the data set. In addition, we exclude layer Va from the analysis because layer Va is not very easy to discriminate in some locations and more importantly it has been shown in previous studies that the response properties to external stimuli in layer Va can be significantly different from layers Vb [17], indicating their functional difference in the process of learning. Last but not least, layer II and layer III may also have functional difference, but it is not easy to distinguish between them basing on the cytoarchitecture architecture [18], so that we had better to analyse them as one layer compartment for a compromise.

In any case, the comparison between cross-location predictions in layers II/III and layer Vb has already revealed the layer-specific long-term memory-dependent dynamics of cortical neural activities. The difference is not due to the relatively different data qualities at different scanning deaths, as we show in the intra-location prediction, the difference between those two layer compartments is much less significant.

In layers II/III, the prediction performances are always quite good in any pair of training-test locations (in the example shown in Figs. 2 A-C, 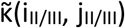 mainly distribute between 0.6 and 0.8; in comparison, in the intra-location prediction in layer II/III of this mouse, 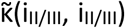 all approximate to 0.8, although technically they cannot be comparable due to the different sizes of training, validation and test data sets). It means in layers II/III, the cortical memory presentations have very similar long-term dynamics across cortical areas. This results is not equal to, but matches the results in previous studies that memory trace neurons were found in layer II/III, insensitive to the cortical areas [9]. The difference is in our research, we do not focus on memory trace cells, but the whole pattern of neural activities. Further analysis reveals that the neural activity patterns in layers II/III are always sensitive to the very recent activities in history, which implies ongoing dynamics in layers II/III with the time scale of one to several days. Its functional role in learning and memory need to study in future research.

On the other hand, in layer Vb, cross-location predictions perform much worse (in the example shown in Figs. 2A-C, some 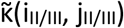 can be as low as 0.2; in comparison, in the intra-location prediction in layer Vb of this mouse, most of 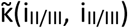 also approximate to 0.8, although again technically they cannot be comparable due to the different sizes of data sets), but between the locations that belong to the same types of cortical areas and same hemispheres, the performances are not too bad, which already implies the different functional roles of cortical areas in layer Vb in the long-term learning and memory process. Consistently, the cross-location predictions within the associative cortices, including posterior parietal cortex and retrosplenial cortex (dorsal) show much more similar performance between layers II/III and layer Vb, whereas the sensory cortices, including visual cortex and somatosensory cortex, show larger difference between those two layer compartments. Results from comparison between the different prediction power distributions further indicate that those information encoded in the neural activities and useful for the neural response to the present environment is segregated and stored in layer Vb in different cortical locations. In other words, when the animal is located in a particular environment, its layer Vb neurons form the patterns as a result of both the response to external multi-modal inputs and the retrieval of previously stored information of different modalities, and obviously the information stored at different previous time points are distributed at different cortical areas. However, we should mention that our approach used in this work is unable to localize the cortical areas for any particular feature of information, which will be an important task in future studies.

### Repeat environment prediction

At the current stage, we could reasonably hypothesize that neural activities in layers II/III are more sensitive to temporal information, but relatively more insensitive to the complexity in terms of sensory modality of the environments compared to layer Vb, whereas when the environment becomes more complex, neural activities in layer Vb will coordinate stronger across cortical areas to perceive the environment. This hypothesis motivated us to test the repeat environment prediction.

Since δ_r_(i_∧_) measures in location i_∧_, how well the present neural activities can predict their activities in a repeated environment in the future, it basically reflects how reliably an environment-specific cortical pattern can be reactivated. Therefore, the variable 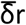 reflects the overall reliability of a layer compartment for reactivating the environment-specific cortical pattern, and σ(δr) reflects the difference of these reliabilities across cortical locations/areas.

It turns out that 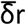 is sensitive to the environment-repeat interval *Inv*, which is consistent with the decay process of memory. In comparison, for layers II/III, 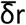 is more sensitive to *Inv* than layer Vb, which verifies the first part of our hypothesis that layers II/III has a more important role in perceiving temporal information than layer Vb.

Among all three pairs of environments we compared, only Training C comprises significantly more sensory modalities than Tunnel, so we expect σ(δr) is smaller in Training C than Tunnel in layers Vb, which turns out to be true (Figs. 3Bd and 3Bf). This result, therefore, verifies the second part of our hypothesis, that layer Vb is more sensitive to the complexity or remembered contexts in terms of sensory modalities.

More interestingly, we know that in Context A and Retrieval B, the animal has significantly different behaviours, that it shows freezing in Retrieval B but not in Context A [9], but the environments Context A and Retrieval B comprise the same sensory modalities. In comparison, their σ(δr) in layer Vb or layers II/III do not have any significant difference (Figs. 3Cd, 3Cf, 3Dd and 3Df). Therefore, the difference in the behaviours is not related to the same aspect of the cortical activities which relates to the sensory modalities of the environments. The only difference of 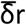 in layers II/III is in fact due to the different environment-repeat intervals (compared Fig. 3Cc to Fig. 3 Ce).

The data of animal Md gives some unexpected results (the last two rows of Fig. 3), but since the studied cortical locations of this animal are limited (only three locations from two cortical areas), it is in difficult to interpret them in a convincing way.

### Regarding the methodology

LightGBM is a machine learning package basing on decision trees. Therefore, its prediction ability is derived from the correlations between the target and the features given that the data are cut into leaves. Similar results could potentially be achieved by correlating the activities on different days. Given the massive number of data points, it is also possible that some deep learning methods might give better prediction results than LightGBM. However, a higher prediction accuracy was not our goal in this work, and deep learning methods usually cannot reveal the deeper mechanisms underlying the different dynamics as revealed here based on the prediction power distributions.

## Conclusions

Activities of cortical neurons are sensitive to both the present environment in which the animals receive stimuli from various modalities as well as the history of past activities reflecting the learned experience of various types of environments, forming long-term memory-dependent activation dynamics. These long-term dynamics are specific for different cortical layers. In layers II/III, they are similar across different cortical areas and different hemispheres, implicating a distributed cortical memory system in layer II/III that integrates multisensory information into the memory. The layer II/III memory network shows ongoing dynamics with a time scale of one to several days. In layer Vb, such consistent memory signal dynamics across-time were lost and their patterns were varied among cortical locations. Between the locations that belong to different types of cortical areas, or belong to different hemispheres, the differences between the long-term memory-dependent dynamics tend to be bigger. Thus, information that has been stored at different previous time points is distributed across layers Vb of different cortical areas, which determine the present activity patterns, jointly with the current multimodal inputs from the environment. Different roles of layer II/III and layer Vb neurons in cross-modal learning process are, therefore, suggested by the layer-specific long-term dynamics of cortical memory representations.

## Data Availability

The datasets generated for this study are available upon request to (J.-S. G.).

## AUTHOR CONTRIBUTIONS

J.-S. G. and C. C. H. designed the research; G. W., H. X., and Y. H. worked on the experiments and collected the data; D. L., and G. W. analysed the data; D. L., J.-S. G., and C. C. H. wrote the paper; all authors approved the paper.

## ACKNOWLEDGMENTS

The authors thank Changsong Zhou for helpful discussions.

